# Two new yeast species of the genus *Vishniacozyma* isolated from the phylloplane of *Equisetum sylvaticum*

**DOI:** 10.1101/2025.02.18.638837

**Authors:** Wladyslav Golubev

## Abstract

Five yeast strains belonging to the genus *Vishniacozyma* Liu et al. were isolated from horsetail in Moscow region (Russia). By morphological and physiological properties, they are similar to the species *V. carnescens, V. peneaus* and *V. victoriae* but differ in mycocin-sensitivity profiles. These isolates represent two novel species, for which the names *V. equsetis* (type strain VKM Y-2979) and *V. paraequisetis* (type strain VKM Y-2980) are proposed.

## Introduction

It is difficult to point out the reviews, dealing with yeast occurrence in nature, that did not report about isolations of *Cryptococcus laurentii* (Kufferath) Skinner (= *Papiliotrema laurentii* Liu et al.) from various sources and substrates. As a rule, the global distribution is typical of taxonomically heterogeneous species that are difficult to distinguish by conventional methods. Actually, the heterogeneity of *P. laurentii* has been recognized in molecular biochemical studies (Sugita et al., 2000, Takashima et al., 2003). However, this assemblage of species (which by their properties fit the standard description of *P. laurentii*) is not yet disclosed fully. By this time some more such species were described (Golubev et al., 2003, 2006, 2008). Initially they were revealed by mycocinotyping and then subsequent closer inspection of isolates led to the conclusion that they represented unknown species. These studies have shown that mycocinotyping (Golubev, 2012) is effective method for distinction of phenotypically similar species.

In the course of a study of epiphytic yeasts from pteridophyte plants (Golubev, 2008) rather many *Cryptococcus* isolates could be identified as *Cr. laurentii* but mycocinotyping showed that they differed both from the type strain of this species and other members of the *Cr. laurentii* complex. This paper deals with characterisation of those isolates and with the formal descriptions of two new species.

## Materials and methods

### Strains

The strains (isolation numbers: Eq-10 = VKM Y-2979, Eq-14, Eq-21 = VKM Y-2980, Eq-26 and Eq-36 = VKM Y-2981) were isolated from the phylloplane of *Equisetum sylvaticum* L. collected in a mixed forest in June. Other strains used in this study were from the Russian Collection of Microorganisms.

### Yeast isolation

Serial dilutions of leaf washings were plated onto infusion agar and incubated at room temperature for 2 weeks. The total culturable yeast populations on *E. sylvaticum* were 20-130·10^3^ cells g^-1^. Infusion agar was prepared in the following way: 200 g of *E. sylvaticum* were sliced into pieces, put into flasks with 1 liter of water and boiled for 10 min. The liquid was filtered through gauze, and water was added to the filtrate to make up 1 liter. After adding 20 g of agar, the medium was autoclaved for 15 min at 15 lbs overpressure. When the temperature cooled down to 50-60°C, streptomycin (500 mg) was added, and the medium was poured into sterile Petri dishes.

### Yeast characterisation

Standard methods in yeast taxonomy were employed for morphological and physiological characterisation (Yarrow, 1998). The procedure for determining mycocin sensitivity patterns was described previously (Golubev et al., 2006).

## Results and Discussion

The following properties of the isolates placed them originally in the genus *Cryptococcus* Vuillemin: lack of sexual state, asexual reproduction by budding, ballistoconidia and arthroconidia not produced, no fermentation, positive urease test, assimilation of *i*-inositol and D-glucuronate, production of starch-like polysaccharides. These strains are nitrate-negative, utilize sucrose, maltose, lactose and melibiose (Fonseca et al., 2011). According to their physiological profiles, they are similar to the species of the *Cr. laurentii* complex. However, in contrast to *Cr. laurentii* type strain our isolates were resistant to mycocins produced by *T. nemorosus* VKM Y-2906, *Saitozyma podzolica* VKM Y-2249 and *Filobasidium capsuligenum* VKM Y-1439. In this respect they were similar to other members of this complex: *Cr. carnescens, Cr. peneaus* and *Cr. victoriae*. At present, these species were assigned to separate genus *Vishniacozyma* on base molecular sequence data (Liu et al.).

Two (Eq-14 and Eq-21) of these isolates are sensitive to *Cr. perniciosus* mycocin, grow at 25°C and assimilate lactose strongly but not L-sorbose as against three ones (Eq-10, Eq-26 and Eq-36). Also, these groups differ in the rate of utilization of some carbon sources.

Sequencing of the D1/D2 domain of the 26S rDNA confirmed that the isolates from *E. sylvaticum* belong to *Cr. laurentii* complex. The closest species to the strain VKM Y-2980 (=Eq-21) was *V. victoriae* but nine nucleotide mismatches were found between them. The strains VKM Y-2979 (= Eq-10) and VKM Y-2981 (= Eq-36) were identical in the D1/D2 region. More discrepancies were found in the ITS region between the strains under study. So, from 22 to 27 differences and by mycocin sensitivity patterns (Table) were observed between the strain VKM Y-2980 and *V. carnescens, V. victoriae*, and from 9 to 17 differences between the strains VKM Y-2979, VKM Y-2981 and *P. laurentii* (J.P. Sampaio, personal communication).

**Table.**
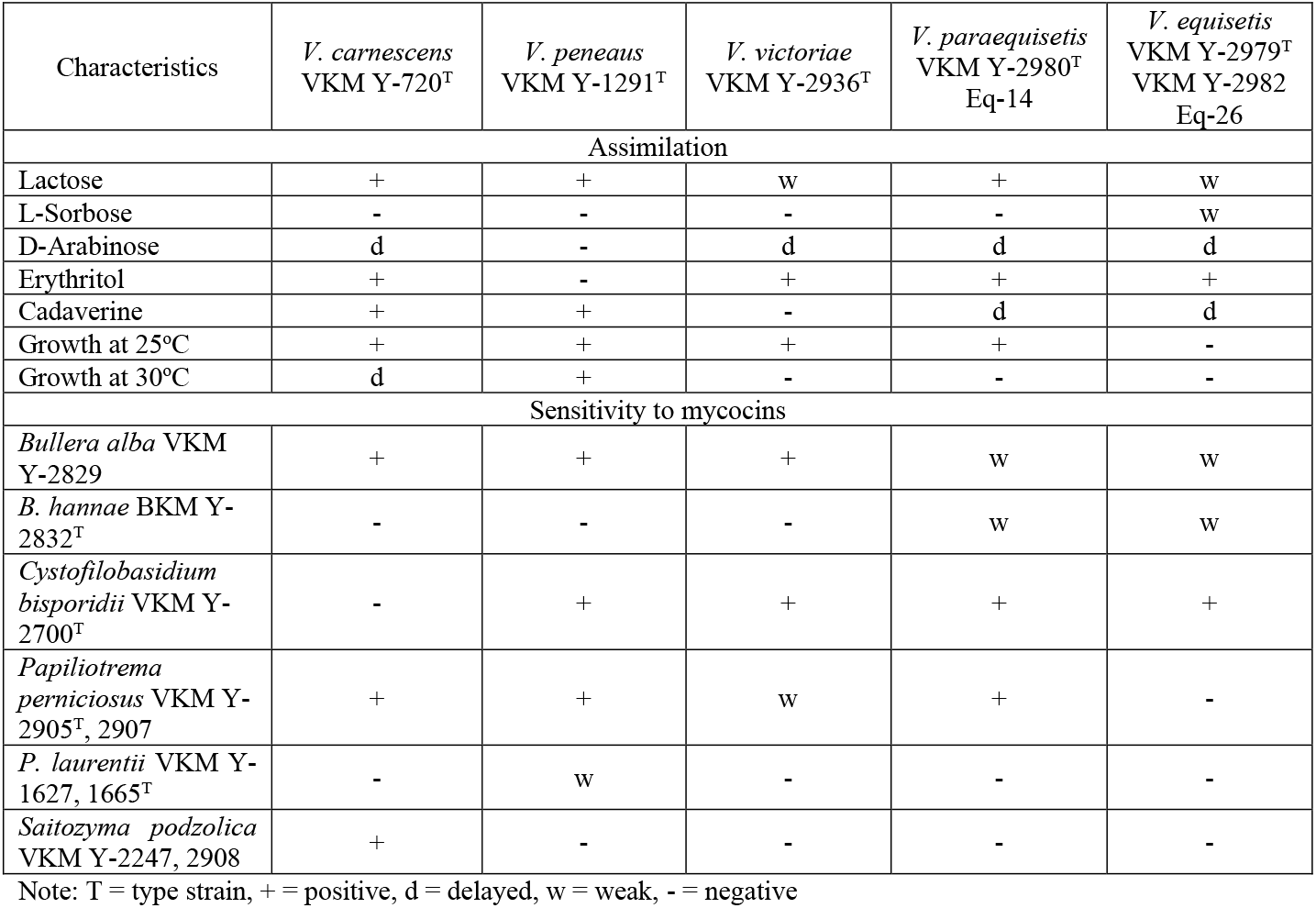
Characteristics distinguishing phylogenetically related *Vishniacozyma* species.

Both analyses showed that our isolates were not conspecific with the closest phylogenetic relatives but hold the separate taxonomic positions. Consequently, the new species, *Vishniacozyma equisetis* and *Vishniacozyma paraequisetis*, are proposed. The specific epithets refer to the origin of the isolates.

The new species assimilate almost all carbon compounds listed in the standard descriptions of species and their physiological characteristics are rather similar. However, they can be distinguished by the combination of some assimilation tests.

### Description of *Vishniacozyma equisetis* Golubev sp. nov

After 3 days in glucose-yeast extract-peptone broth, cells are subglobose and oval [width/length ratio 1.2-1.8 (mean 1.4 μm)], 3.4-4.3 × 4.2-6.0 μm (mean 3.7 × 5.2 μm) with capsules and single or in pairs. After 1 month, there is a sediment, a ring and islets. After 1 month, the streak culture on yeast morphology agar (Difco) is cream, smooth, glistening with an entire margin. No ballistoconidia are observed. After 10 days in slide cultures on cornmeal agar, neither true mycelium nor pseudomycelium is produced. Fermentation is absent. The following carbon compounds are assimilated: D-glucose, galactose, L-sorbose (weak), α-methyl-D-glucoside, D-ribose (slow), D-xylose, L-arabinose, D-arabinose (slow), L-rhamnose, D-glucosamine (slow), N-acetyl-D-glucosamine, sucrose, maltose, trehalose (slow), cellobiose, lactose (weak), melibiose, melezitose, raffinose, arbutin, salicin (slow), soluble starch (slow), glycerol (weak), erythritol, xylitol, L-arabitol, sorbitol, mannitol (slow), galactitol, i-inositol, D-glucuronate, gluconate, 2-keto-D-gluconate, 5-keto-D-gluconate, glucarate (slow), succinate, citrate (slow) and lactate (slow). No growth occurs on inulin, ethanol, quinic acid, glycine, orcinol and allantoin. Utilization of nitrogen compounds: positive for L-lysine, ethylamine (slow), D-glucosamine and cadaverine (slow), negative for nitrate, nitrite, creatine and creatinine. Urease and starch-like compounds are produced. Growth in vitamin-free medium is absent. Growth at 25°C is negative. Strains are *bisporidii* VKM Y-2700, but are resistant to the mycocins of *B. sinensis* var. *lactis* VKM Y-2826, *B. unica* VKM Y-2830, *laurentii* VKM Y-1627, VKM Y-1628, VKM Y-1665, *Papiliotrema nemorosus* VKM Y-2906, *P. perniciosus* VKM Y-2905, VKM Y-2907, *Saitozyma. podzolica* VKM Y-2247, VKM Y-2249, VKM Y-2908, *Cyst. infirmominiatum* VKM Y-2897 and *Filobasidium capsuligenum* VKM Y-1439. The type strain, Eq-10 (VKM Y-2979), was isolated from *Equisetum sylvaticum* L. in the Moscow region (Russia) and has been deposited in the Russian Collection of Micro-organisms (Pushchino, Russia). GenBank accession numbers: D1/D2 - HM749316, ITS - HM749321.

### Description of *Vishnicozyma paraequisetis* Golubev sp. nov

After 3 days in glucose-yeast extract-peptone broth, cells are subglobose and oval [width/length ratio 1.1 - 1.4 (mean 1.2 μm)], 3.4-6.0 × 4.2-6.8 μm (mean 4.4 × 5.4 μm) with capsules and single or in pairs. After 1 month, there is a sediment and a pellicle. After 1 month, the streak culture on yeast morphology agar (Difco) is cream, smooth and glistening with an entire margin. No ballistoconidia are observed. After 10 days in slide cultures on cornmeal agar a rudimentary pseudomycelium is produced. Fermentation is absent. The following carbon compounds are assimilated: D-glucose, galactose, α-methyl-D-glucoside, D-ribose (slow), D-xylose, L-arabinose, D-arabinose (slow), L-rhamnose, D-glucosamine (slow), N-acetyl-D-glucosamine, sucrose, maltose, trehalose (weak), cellobiose, lactose, melibiose, melezitose, raffinose, arbutin, salicin (slow), inulin (weak), soluble starch (slow), ethanol (weak), glycerol (slow), erythritol, xylitol, L-arabitol, sorbitol, mannitol, galactitol (slow), i-inositol, D-glucuronate, gluconate (weak), 2-keto-D-gluconate, 5-keto-D-gluconate, glucarate (weak), succinate (slow), citrate (weak) and lactate (weak). No growth occurs on L-sorbose, quinic acid, glycine, orcinol and allantoin. Utilization of nitrogen compounds: positive for L-lysine, ethylamine (slow), D-glucosamine and cadaverine (slow), negative for nitrate, nitrite, creatine and creatinine. Urease and starch-like compounds are produced. Growth in vitamin-free medium is absent. Grows at 25°C but not at 30°C. Strains are sensitive to the mycocins produced by *Bullera alba* VKM Y-2829, *B. hannae* VKM Y-2832, *Papiliotrema perniciosus* VKM Y-2905, VKM Y-2907 and *Cystofilobasidium bisporidii* VKM Y-2700, but are resistant to the mycocins of *B. sinensis* var. *lactis* VKM Y-2826, *B. unica* VKM Y-2830, *P. laurentii* VKM Y-1627, VKM Y-1628, VKM Y-1665, *P. nemorosus* VKM Y-2906, *Saitozyma podzolica* VKM Y-2247, VKM Y-2249, VKM Y-2908, *Cyst. infirmominiatum* VKM Y-2897 and *Filobasidium. capsuligenum* VKM Y-1439. The type strain, Eq-21 (VKM Y-2980), was isolated from *Equisetum sylvaticum* L. in the Moscow region (Russia) and has been deposited in the Russian Collection of Micro-organisms (Pushchino, Russia). GenBank accession numbers: D1/D2 - HM749318, ITS - HM 749319.

## Acknowledgemets

The author is grateful to J.P. Sampaio (Univ. Nova de Lisboa, Portugal) for the molecular biological data. This work was supported by Ministry of Science and Higher Education of the Russian Federation, agreement no. 075-15-2021-1051.

## Funding

### Statements and Declarations

This work was supported by Ministry of Science and Higher Education of the Russian Federation, agreement no. 075-15-2021-1051.

## References

Fonseca A., Boekhout T., Fell J.W. Cryptococcus Vuillemin (1901). In: The Yeasts, a Taxonomic Study. 5th ed. (Kurtzman C.P., Fell J.W., Boekhout T., eds.). 2011. V. 3. pp. 1661–1737.

Golubev W.I. Epiphytic yeasts from pteridophyte plants. Abstr. 2nd Congress of Russian Mycologists (16-18th April 2008, Moscow). pp. 272–273 (In. Russ.).

Golubev W.I., Gadanho V., Sampaio J.P., Golubev N.W. (2003) Cryptococcus nemorosus sp. nov. and Cryptococcus perniciosus sp. nov., related to Papiliotrema Sampaio et al. (Tremellales). Int. J. System. Evol. Microbiol. V. 53, pp. 905–911. 10.1099/ijs.0.02374-0

Golubev W.I., Sampaio J.P., Alves L., Golubeva E.W. (2006) Cryptococcus silvicola nov. sp. from nature reserves of Russia and Portugal. Ant. van Leeuwenh. V. 89, pp. 45–51. 10.1007/s10482-005-9008-z

Golubev W.I., Pfeiffer I., Tomashevskaya M.A. (2008) Cryptococcus pinus sp. nov., an anamorphic basidiomycetous yeast isolated from pine litter. Int. J. System. Evol. V. 68, pp. 1968–1971. 10.1099/ijs.0.65764-0

Golubev W.I. (2012) Mycocinotyping. Mycology and Phytopathology. V. 46, pp. 3–13 (In Russ.).

Liu X.Z., Wang Q.M., Groenwald M. et al. (2015) Towards an integrated phylogenetic classification of the Tremellomycetes. Studies in Mycology. V. 81, pp. 85–147. 10.1016/j.simyco.2015.12.001

Sugita T., Takashima M., Ikeda R., Nakase T., Shinoda T. (2000) Intraspecies diversity of Cryptococcus laurentii as revealed by sequences of internal transcribed spacer regions and 28S rRNA gene and taxonomic position of C. laurentii clinical isolates. J. Clin. Microbiol. V. 38, pp. 1468–1471. 10.1128/jcm.38.4.1468-1471.2000

Takashima M., Sugita T., Shinoda T., Nakase T. (2003) Three new combinations from the Cryptococcus laurentii complex: Cryptococcus aureus, Cryptococcus carnescens and Cryptococcus peneaus. Int. J. System. Evol. Microbiol. V. 53, pp. 1187–1194. 10.1099/ijs.0.02498-0

Yarrow D. (1998) Methods for isolation, maintenance and identification of yeasts. In: The Yeasts. A Taxonomic Study. 4th ed. (Kurtzman C.P., Fell J.W., eds.) Elsevier. Amsterdam. pp. 77–100.

